# Chimeric mRNA based COVID-19 vaccine induces protective immunity against Omicron and Delta

**DOI:** 10.1101/2022.03.04.483032

**Authors:** Qidong Hu, Ying Zhao, Namir Shaabani, Xiaoxuan Lyu, Haotian Sun, Vincent Cruz, Yi Kao, Jia Xu, Amber Fossier, Karen Stegman, Zhihao Wang, Zhenping Wang, Yue Hu, Yi Zheng, Lilian Kyaw, Cipriano Zuluaga, Hua Wang, Hong Pei, Colin Powers, Robert Allen, Hui Xie, Henry Ji, Runqiang Chen

**Affiliations:** Sorrento Therapeutics Inc., 4955 Directors Place, San Diego, CA, USA

## Abstract

The emerging SARS-CoV-2 variants of concern (VOCs) exhibit enhanced transmission and immune escape, reducing the efficacy and effectiveness of the two FDA-approved mRNA vaccines. Here, we explored various strategies to develop novel mRNAs vaccines to achieve safer and wider coverage of VOCs. Firstly, we constructed a cohort of mRNAs that feature a furin cleavage mutation in the spike (S) protein of predominant VOCs, including Alpha (B.1.1.7), Beta (B.1.351), Gamma (P.1) and Delta (B.1.617.2). Not present in the mRNA vaccines currently in use, the mutation abolished the cleavage between the S1 and S2 subunits, potentially enhancing the safety profile of the immunogen. Secondly, we systematically evaluated the induction of neutralizing antibodies (nAb) in vaccinated mice, and discovered that individual VOC mRNAs elicited strong neutralizing activity in a VOC-specific manner. Thirdly, the IgG produced in mice immunized with Beta-Furin and Washington (WA)-Furin mRNAs showed potent cross-reactivity with other VOCs, which was further corroborated by challenging vaccinated mice with the live virus of VOCs. However, neither WA-Furin nor Beta-Furin mRNA elicited strong neutralizing activity against the Omicron variant. Hence, we further developed an Omicron-specific mRNA vaccine that restored protection against the original and the sublineages of Omicron variant. Finally, to broaden the protection spectrum of the new Omicron mRNA vaccine, we tested the concept of bivalent immunogen. Instead of just fusing two RBDs head-to-tail, we for the first time constructed an mRNA-based chimeric immunogen by introducing the RBD of Delta variant into the entire S antigen of Omicron. The resultant chimeric mRNA was capable of inducing potent and broadly acting nAb against Omicron (both BA.1 and BA.2) and Delta, which paves the way to develop new vaccine candidate to target emerging variants in the future.

## Introduction

The global pandemic caused by the severe acute respiratory syndrome coronavirus 2 (SARS-CoV-2) has already claimed millions of lives since the outbreak in January 2020^1^. Numerous strategies have been employed to combat the spread of this infectious disease by public health agencies worldwide. As one of the most widely adopted approaches, vaccination has been proved to be a key tool to tackling COVID-19. To date, there are a total of ten COVID-19 vaccines granted for emergency use or fully approved globally^2^. Among the various types of vaccine platforms, the mRNA vaccine has been proved to be effective against COVID-related hospitalization and death (https://www.cdc.gov/coronavirus/2019-ncov/science/science-briefs/fully-vaccinated-people.html).

Despite these efforts, the pandemic continues to pose a threat to the public health due to the constant viral evolution and the consequential emergence of new SARS-CoV-2 variants of concern (VOCs)^3^. Hence, there is an urgent need to develop more efficacious mRNA vaccines to fight against the current and potentially future VOCs.

The designated VOCs have caused multiple waves of widespread outbreak with unprecedented speed. Back in 2020, the first VOC Alpha (B.1.1.7) was detected in the United Kingdom, leading to more than 100,000 confirmed cases in the country. Following the emerging of Alpha variant, there were two other outbreaks originating from South Africa and Brazil, triggered by the Beta (B.1.351) and Gamma (P.1) variants, respectively^1,3,4^. With additional mutation in spike protein, both VOCs were estimated to be 40% −80% more transmissible than the wildtype lineage. Furthermore, the Beta variant exhibits great immune escape and COVID-19 vaccination only showed 75% effectiveness against infection^5^. In late 2020, a new variant B. 1.617.2 (Delta) was identified and subsequentially contributed to a surge in cases in India and worldwide shortly afterwards. Besides the increased transmissibility, Delta variant has been shown to cause more severe disease and result in a poorer prognosis than previously reported VOCs^6–8^. To make things even worse, the effectiveness of existing mRNA vaccines against Delta has dropped considerably, from ~95% for the original WA strain to 76% (mRNA-1273) and 42% (BNT162b2)^9^. By June 2021, global COVID-19-related deaths hit 5 million as Delta variant swept over 161 countries around the world.

More recently, WHO declared a novel VOC B.1.1.529, designated as Omicron (BA.1), on 26^th^ Nov 2021, only two days after it was first reported to WHO by South Africa^10^. Omicron quickly outcompeted the circulating Delta variant within weeks after landing in Europe and US, and has become the dominant VOC around the globe. Scientists found that there was a significant reduction in Omicron virus neutralizing activity in sera obtained from individuals infected by pre-Omicron variants and populations vaccinated with immunogens based on early WA-1 virus^11–13^. A huge number of breakthrough infections have been reported even from individuals fully vaccinated with the 3^rd^ booster shot. Although still largely effective in preventing hospitalization and mortality, the two existing mRNA-based vaccines have shown declining capabilities in preventing infection by various VOCs, especially Omicron. A plausible explanation for the increasingly large gap in vaccine coverage is that both EUA-approved mRNA vaccines encode the S protein of the original WA strain as the immunogen.

Compared to its ancestral strain, Omicron contains more than 30 additional mutations within the S protein coding sequence, 15 of which reside in the receptor binding domain (RBD)^14^. Capable of binding to the human ACE2 receptor, RBD has been demonstrated by pioneering structural analysis to be the key domain for virus attachment and entry to human cells^15^. Thus, the high rate of mutations in Omicron RBD may dramatically alter the interaction dynamics between the virus and host cell, which could, at least partially, explain the enhanced transmissibility and breakthrough cases^4,13^.

In the present study, we explored various strategies to develop new mRNA vaccines that can provide broad and effective protection against predominant SARS-CoV-2 VOCs. We first screened five monovalent mRNA vaccines encoding the spike (S) protein of Alpha, Beta, Gamma, Delta, and the original Washington (WA) strain. All the vaccines harbor the mutation that abolishes the furin-mediated cleavage between S1 and S2 domains of S protein. It was found that Beta-Furin and WA-Furin mRNAs showed the most potent cross-neutralizing activity against other VOCs, except for Omicron. Hence, we further constructed the mRNA vaccine encoding the S protein of Omicron, which was shown in subsequent experiments to induce the strongest protection against Omicron. However, it did not elicit broad neutralizing capacity against other VOCs, such as Delta. To address the observed limitations of an Omicron-specific mRNA vaccine, we developed a chimeric mRNA by incorporating an extra RBD from Delta variant in the context of the S protein of Omicron. The resultant chimeric mRNA vaccine elicited significantly higher cross-neutralization activity against Delta, while retaining efficacy against Omicron. Our findings thus provide insights into the development of the next generation mRNA vaccine with wider coverage of possible future VOCs of SARS-CoV-2, and the potential to provide vaccine coverage for respiratory pathogens outside of the family *Coronaviridae*.

## Results

### mRNA vaccine harboring furin cleavage mutation elicits strong neutralizing titers and T cell responses

Previously, we have designed a SARS-CoV-2 spike (S) mRNA vaccine that achieves high expression in mammalian cells^16^. This mRNA vaccine encodes the S protein from the Wuhan/Washington (WA) strain and encodes a polybasic furin cleavage site at the junction of S1 and S2 subunits. The feature could affect the stability of spike protein and reduce the pool of antigenic epitopes available to induce cellular and humoral immunity^17^. In addition, cleaved S1 was detected in the blood of immunized subjects receiving existing mRNA vaccine^18^, raising a significant question of whether this free moiety could represent a potential safety issue by mimicking full-length S protein to activate ACE2, and thus trigger some side effects, including myocarditis in healthy subjects. Thus, to further optimize the mRNA vaccine and eliminate these potential safety concerns, the furin cleavage site between the S1 and S2 domains of the spike was mutated (**Figure 1A**) in the currently reported build of our mRNA vaccine candidates.

**Figure 1.**
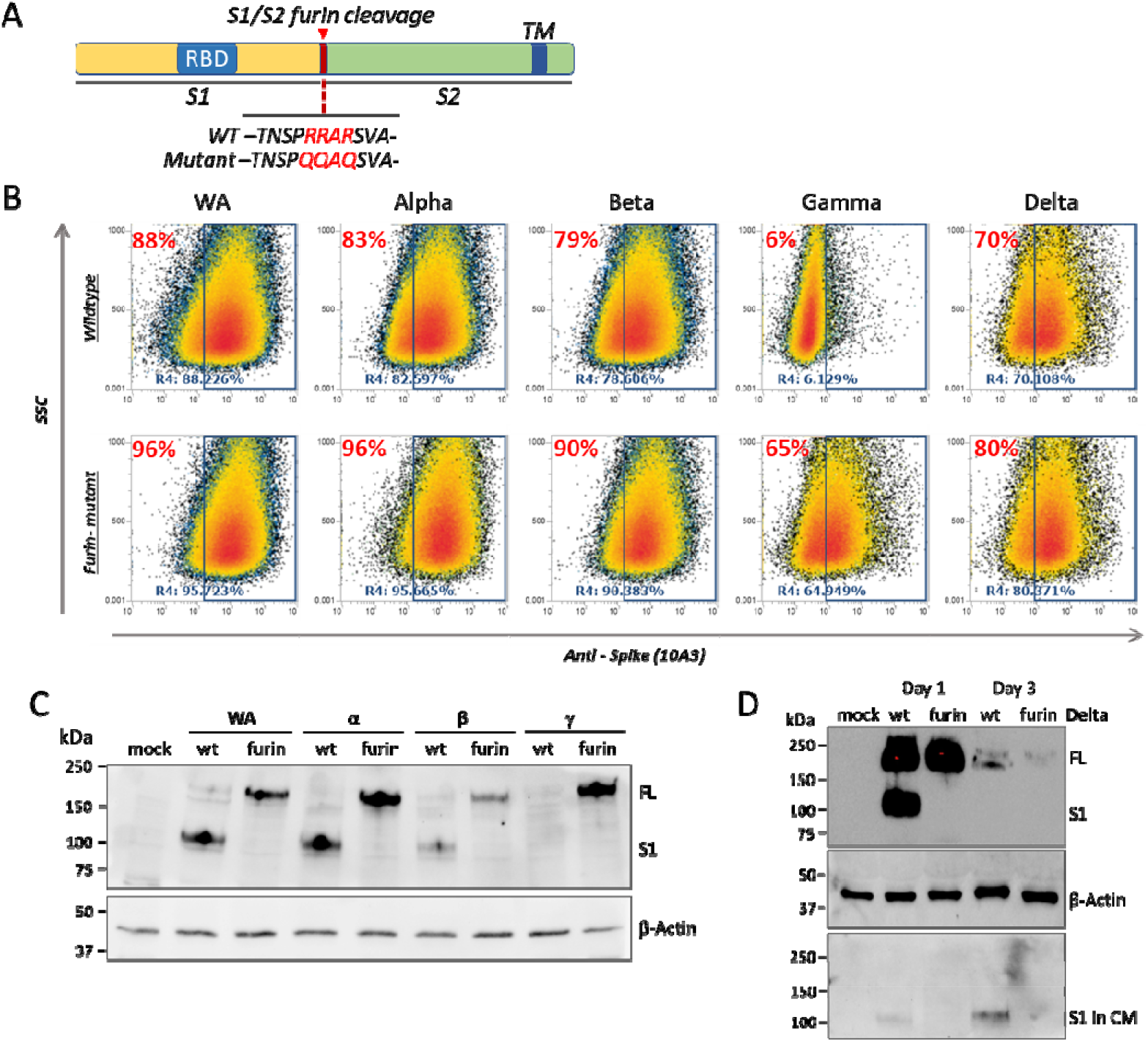
Expression and cleavage of Furin cleavage mutant mRNA. (**A**) Design of the furin cleavage mutant mRNA. (**B**) 293T cells in 24-well plate were transfected with 1 ug indicated variant mRNAs and collected 72 hours post transfection. The flow cytometry using the proprietary anti-spike 10A3 antibody showed moderate increase in the surface expression of spike after furin cleavage mutation. (**C**) 293T cells in 6-well plate were transfected with 5 ug indicated variant mRNAs and collected 24 hours post transfection. Western blot using anti-S1 antibody showed that FL spike was the dominant species after the removal of furin cleavage site. (**D**) 293T cells in 6-well plate were transfected with 5 ug wild-type or furin cleavage mutant spike mRNA of Delta variant. The cell lysate and CM were collected at day 1 and day 3 post transfection. The relative abundance of FL spike and free S1 was examined by Western blot. TM: transmembrane domain; FL: full-length S protein; CM: conditioned medium.

We then constructed two sets of mRNAs encoding the S protein of all the predominant SARS-CoV-2 VOCs, one with the wild-type (WT) furin cleavage site, the other with mutated site. We examined and compared the expression of these VOC mRNAs, including WA, Alpha, Beta, Gamma and Delta in 293T cells. The flow cytometry results showed that removal of the furin cleavage site increased the surface expression of VOC S protein in transfected 293T cells (**Figure 1B**), and Western blot confirmed that the mutation abolished the furin cleavage and lowered the level of free S1 in the conditioned medium (**Figure 1C, D**). We thus hypothesized that the boosted expression of the furin mutant could result in stronger immunogenicity. To test the hypothesis, WA WT or furin mutant mRNAs were formulated with an in-house lipid nanoparticle (LNP), and six-week-old female BALB/c mice were intramuscularly injected with two doses of each LNP-mRNA separated by three weeks. ELISA on the sera collected 14 days after boost revealed that mRNA carrying the furin cleavage mutation elicited a higher average endpoint titer (EPT) of total binding antibody than its WT version (**Figure S1A**). We then performed the Plaque Reduction Neutralization Test (PRNT) using the same antisera to assess neutralizing antibody (nAb)^19,20^. Consistent with ELISA data, PRNT results confirmed the superior neutralizing activity of sera from WA furin mutant-injected mice (**Figure S1B**) as compared to that measured in mice vaccinated with the WT version.

Likewise, to investigate the ability of individual SARS-CoV-2 VOC mRNA vaccines to generate nAb, and to profile their coverage spectrum *in vivo*, we immunized six-week-old female BALB/c mice with LNP-encapsulated furin-mutant VOC mRNAs and compared the performance of individual mRNAs in vaccinated mice. The EPT of total binding antibodies was first measured by ELISA using the sera day 14 post two-dose injection. As expected, each mRNA induced the strongest antibody response against the corresponding VOC S protein, with the exception of Gamma (**Figure 2A**). Intriguingly, some VOC mRNA vaccinations led to the production of antibodies capable of binding to a breadth of VOC S proteins, especially Beta-Furin. To further analyse the neutralizing capability of individual sera, PRNT was performed where VeroE6 cells were exposed to the live virus of five VOCs in the absence or presence of diluted serum collected from the immunized mice (**Figure 2B**). In keeping with the ELISA results, the experiment showed that the individual monovalent mRNA vaccine generally displayed variant-specific protection activity. For example, WA-Furin mRNA induced the highest neutralizing activity against the WA-1 virus. Again it was observed that the serum from the Beta-Furin mRNA injected cohort displayed robust and broad protection against all VOCs tested. Particularly, the Beta-Furin mRNA provided a much stronger protection against the highly contagious Delta variant than was elicited by vaccination with the WA-Furin mRNA. The data indicates that whereas the VOC-specific strategy can provide strain-specific protection, some VOC-based mRNA vaccines have relatively enhanced potential to trigger a broad and potent immune response to the genetically divergent set of existing SARS-CoV-2 variants.

**Figure 2.**
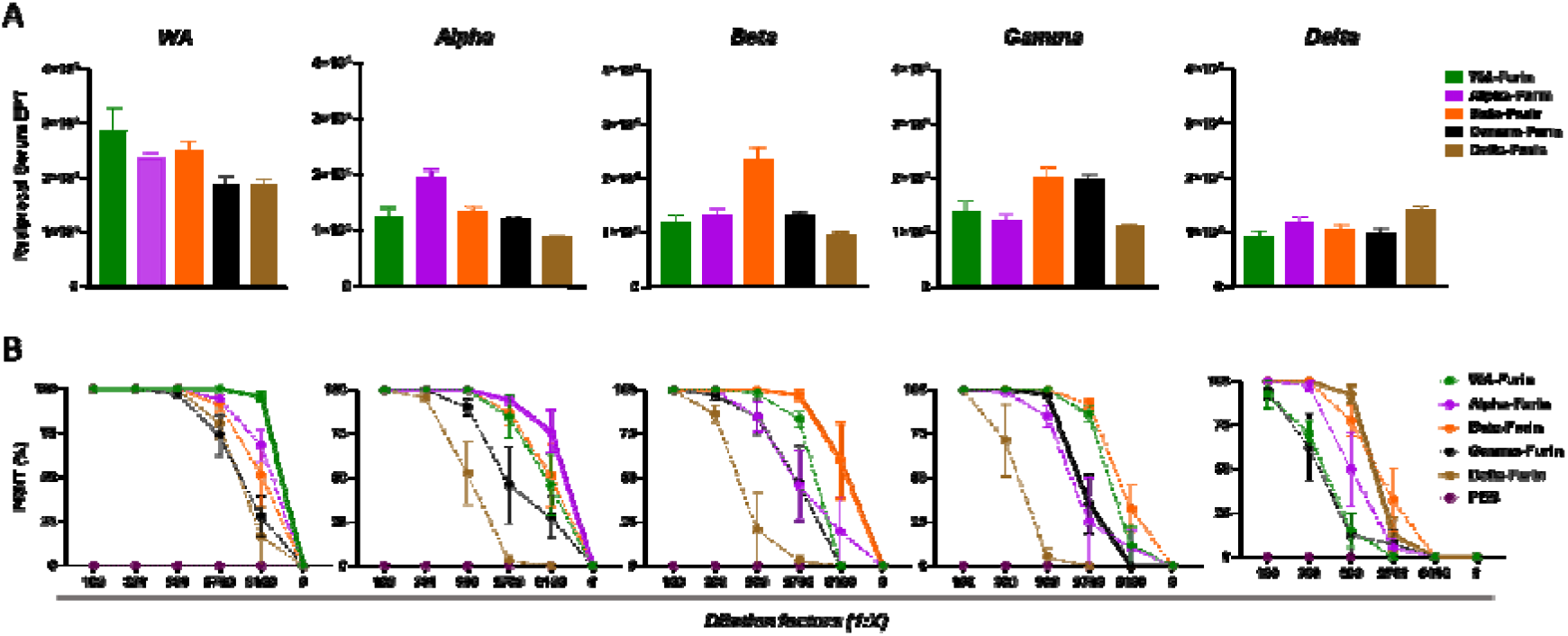
Antibody responses upon immunization with VOC-based vaccines *in vivo*. Six-week-old female BALB/c mice were immunized twice with indicated furin-mutant VOC mRNAs with a 3-week interval. (**A**) Day 14 sera post booster were evaluated for antibodies binding to recombinant S protein from designated VOCs by ELISA. (**B**) The Day 14 sera post booster were evaluated for nAb responses against designated live virus by 50% plaque reduction neutralization test (PRNT). Plots: mean±SEM; n=6. EPT: endpoint titer.

To profile the TH1-biased and TH2-biased response following immunization, Balb/c mice received two injections of WA-Furin or Beta-Furin mRNA at 0.2 ug, 1.0 ug and 5.0 ug. Seven weeks after the booster injection, splenocytes were isolated and stimulated with the peptide pool that covers the S1 subunit of the S protein. Intracellular cytokine staining showed that both mRNAs elicited a dose-dependent TH1 response as exemplified by IFNγ and TNFα, with limited TH2 response indicated by IL-4 and IL-5 (**Figure 3A, B**). The mRNAs also triggered a potent induction of IFNγ and TNFα expression in CD8+ T cells in a dose-dependent manner (**Figure 3A, B**). To further assess the effect of our mRNA vaccines on T cell composition and formation of memory B cells, we quantified CD4+ T cells, CD8+ T cells and memory B cells isolated from spleen, as well as memory B cells from blood. The result showed minimal changes in the percentage of these cells (**Fig. S2**).

**Figure 3.**
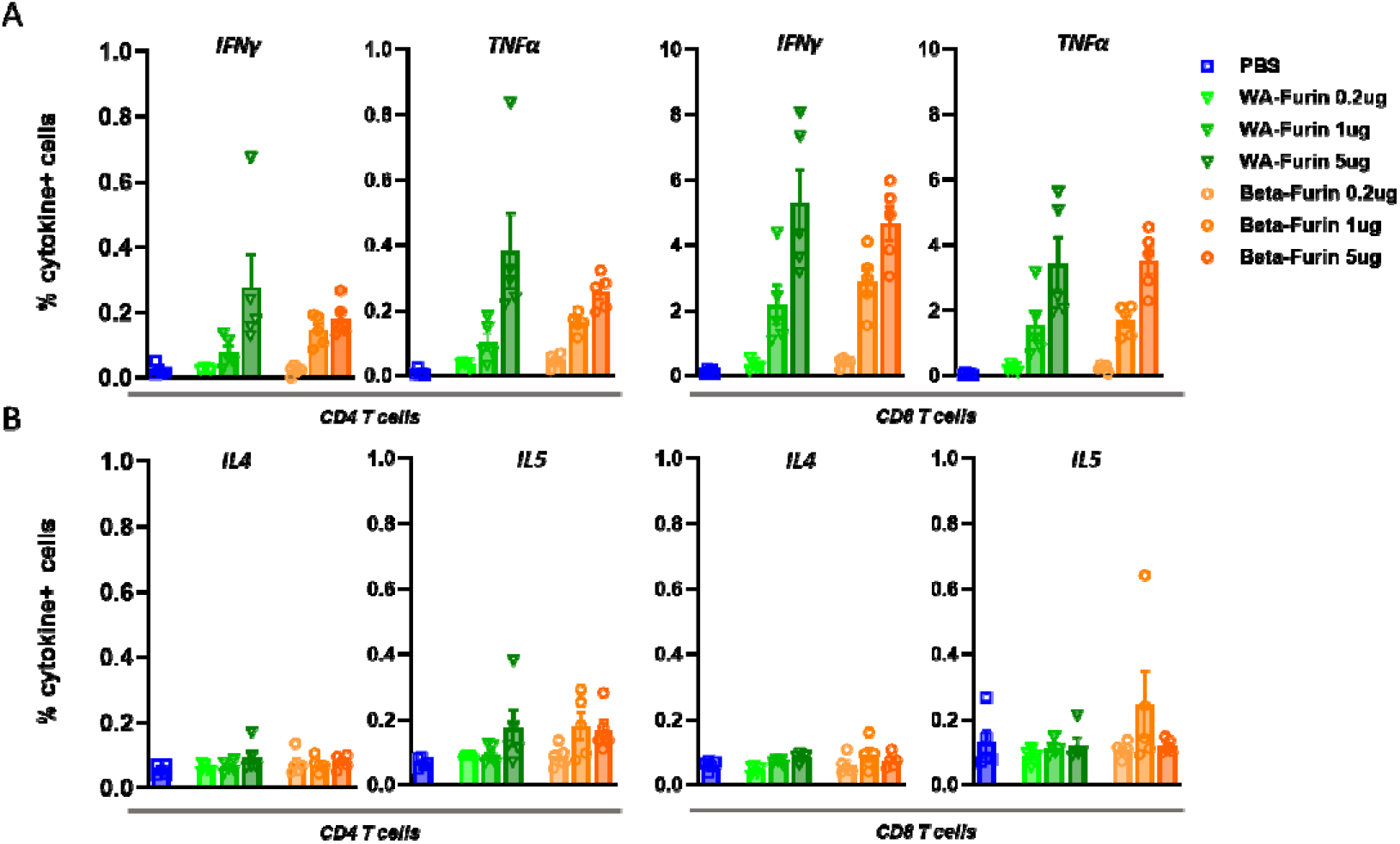
Phenotyping of CD4+ and CD8+ T lymphocytes in spleen of immunized mice. Six-week-old female BALB/c mice were immunized twice with WA-Furin or Beta-Furin mRNA at indicated doses with a 3-week interval. Seven weeks after booster, splenocytes were stimulated *ex vivo* with the S1 peptide pool for 6 hours. Intracellular cytokine staining was then performed to measure TH1-related IFNγ and TNFα (**A**), and TH2-related IL-4 and IL-5 (**B**). Bars: mean±SEM; n=5.

### mRNA vaccines carrying furin cleavage mutation produce robust protection *in vivo*

The K18-hACE2 transgenic model has been extensively utilized to evaluate the vaccine efficacy and effectiveness in preventing COVID-19 in the preclinical setting^21–24^. Two key metrics to determine the severity of pathogenesis are the virus titer in the lung tissue and body weight loss following virus infection. To investigate the protection capacity of our furin-mutant mRNA vaccines, K18-hACE2 mice were first intramuscularly administrated with 5μg of WA-Furin or Beta-Furin mRNA twice with 3-week interval. Five weeks post full vaccination, the animals were challenged with 1 × 10^5^ half-maximal tissue culture infectious dose (TCID50) of the WA, Beta or Lambda variants (**Figure 4A**). Then the virus replication in the lung was quantified to determine the effect of vaccination. The average virus titer was approximated to 1 × 10^6^, 2.1 ×10^5^, and 4.5 × 10^6^ TCID50/g for WA, Beta and Lambda strains, respectively in the control group injected with PBS. On the other hand, immunization with either WA-Furin or Beta-Furin mRNA almost completely inhibited the replication of virus in the lungs, with virus titers falling below the limit of detection (**Figure 4B**). As expected, animals treated with PBS exhibited dramatic weight loss in all challenge settings, regardless of the virus strain. The average body weight in the PBS controls declined to 82%, 78% and 81% on day 5 post infection with WA, Beta and Lambda strains, respectively. In contrast, none of the mice immunized with Beta-Furin or WA-Furin vaccines showed any sign of weight loss, and some animals in these treatment groups gained weight after infection up to 5 days (**Figure 4C**). In addition, both furin-mutant mRNAs gave effective protection against the Lambda variant although the corresponding spike mRNA was not included among the immunogens, suggesting broad protection capacity of some VOC mRNAs.

**Figure 4.**
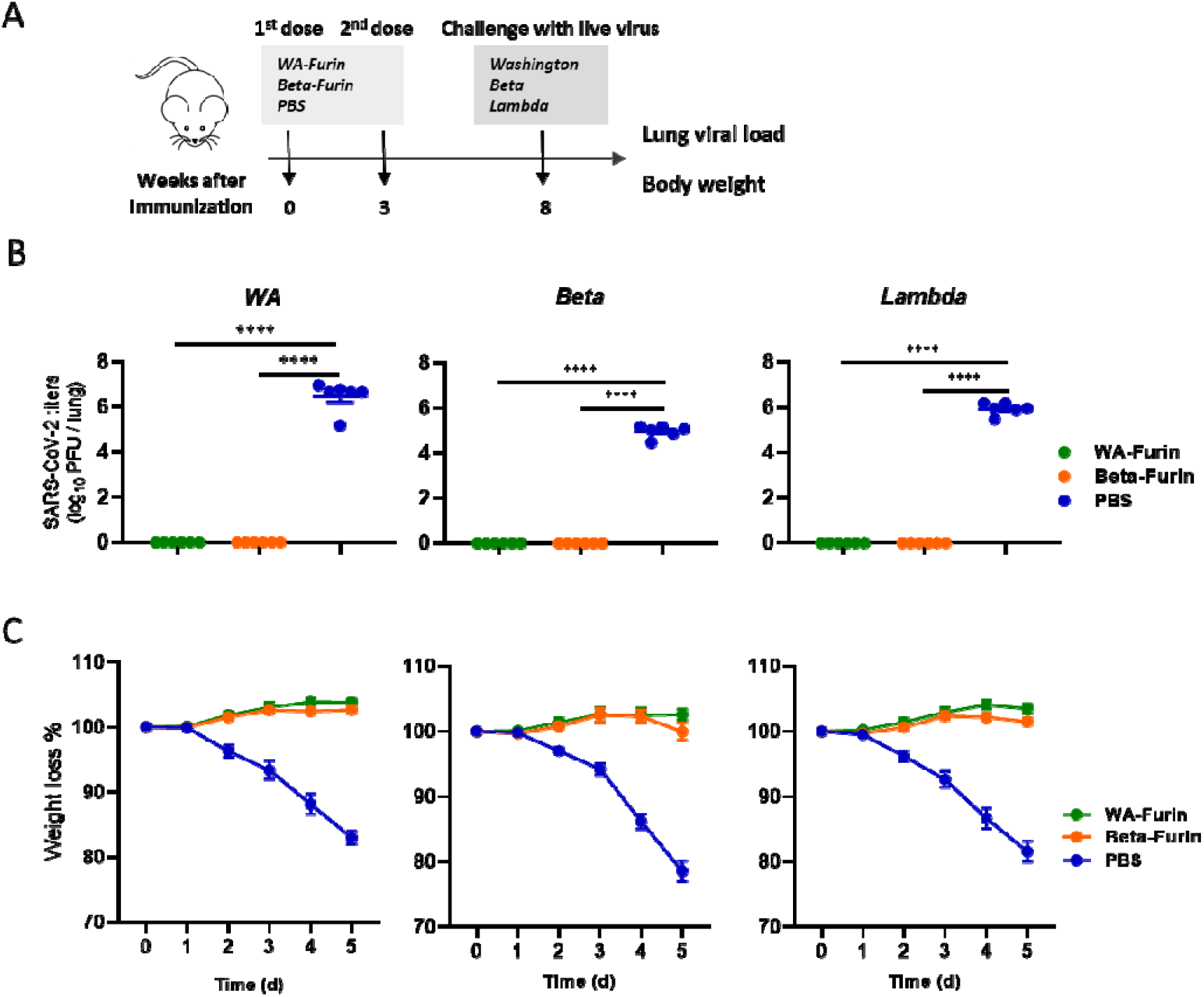
Immunization with Furin cleavage mutant vaccine produces strong protection *in vivo*. (**A**) Schematic illustration of the design of the *in vivo* study using K18 hACE2 mice. The lung virus titer (**B**) and body weight (**C**) were measured in mice challenged with the live virus of indicated SARS-CoV-2 VOCs after full immunization. Plots: mean±SEM; n=6. ****p<0.0001 one-way ANOVA.

### Omicron variant undermines the effectiveness of the past VOC based vaccines

Recently, a new SARS-CoV-2 variant B.1.1.529 (Omicron), was identified in South Africa and declared as VOC by WHO due to its rapid rise in global prevalence. Bearing an unusually large number of previously unreported RBD mutations as compared to other VOCs, Omicron exhibited the most significant escape from the serum of both fully vaccinated subjects and convalescents^25^. To investigate whether our optimized mRNA vaccines could offer effective protection against Omicron, VSV pseudotyped viruses were used to determine the neutralization potency of sera collected from vaccinated animals. In agreement with other studies^26^, we found that the mean neutralization titer (IC50) of Omicron was only 318 for the sera collected at day 14 after boost with WA-Furin mRNA, which represents a significant decrease of neutralization, as compared to those recorded against other VOC-based pseudotyped viruses (**Figure S3**). A similar decrease in the Omicron protection capacity of the sera was observed for mRNA vaccine candidates derived from other VOCs (**Figure S3**). Surprisingly, even the Beta-Furin mRNA, which has been shown to provide the broadest immune response against all major VOCs (**Figure 2**), failed to induce sufficient protection against Omicron, prompting us to explore further immunogen designs to overcome this liability.

To establish the potential of a variant-matched monovalent vaccine to provide protective immunity against Omicron virus, we replaced the coding sequence of the original mRNA vaccines with Omicron spike retaining the furin cleavage mutation. More than 85% of the Omicron-Furin mRNA transfected 293T cells, but not the mock control, could be detected by flow cytometry with Fc-tagged ACE2 recombinant protein (**Figure S4**). We then performed *in vivo* studies to assess the immunogenicity and efficacy of the Omicron-specific mRNA vaccine. BALB/c mice were immunized intramuscularly with 5μg Omicron-LNP mRNA or Beta-LNP mRNA two times at a three-week interval. Serum was collected two weeks post boost and subjected to ELISA assay to measure Omicron-specific binding antibodies. Using the recombinant Omicron spike protein as the coating antigen, high titers of binding antibodies were observed in the sera of Omicron mRNA-injected mice (**Figure 5A**). The protective capability of Omicron-specific mRNA vaccine was further evaluated in PRNT. We confirmed that the sera collected from Omicron-Furin-immunized animals offered superior protection against the infection of Omicron strain (**Figure 5B**), demonstrating that this newly designed mRNA vaccine could induce potent production of Omicron-specific nAbs *in vivo*.

**Figure 5.**
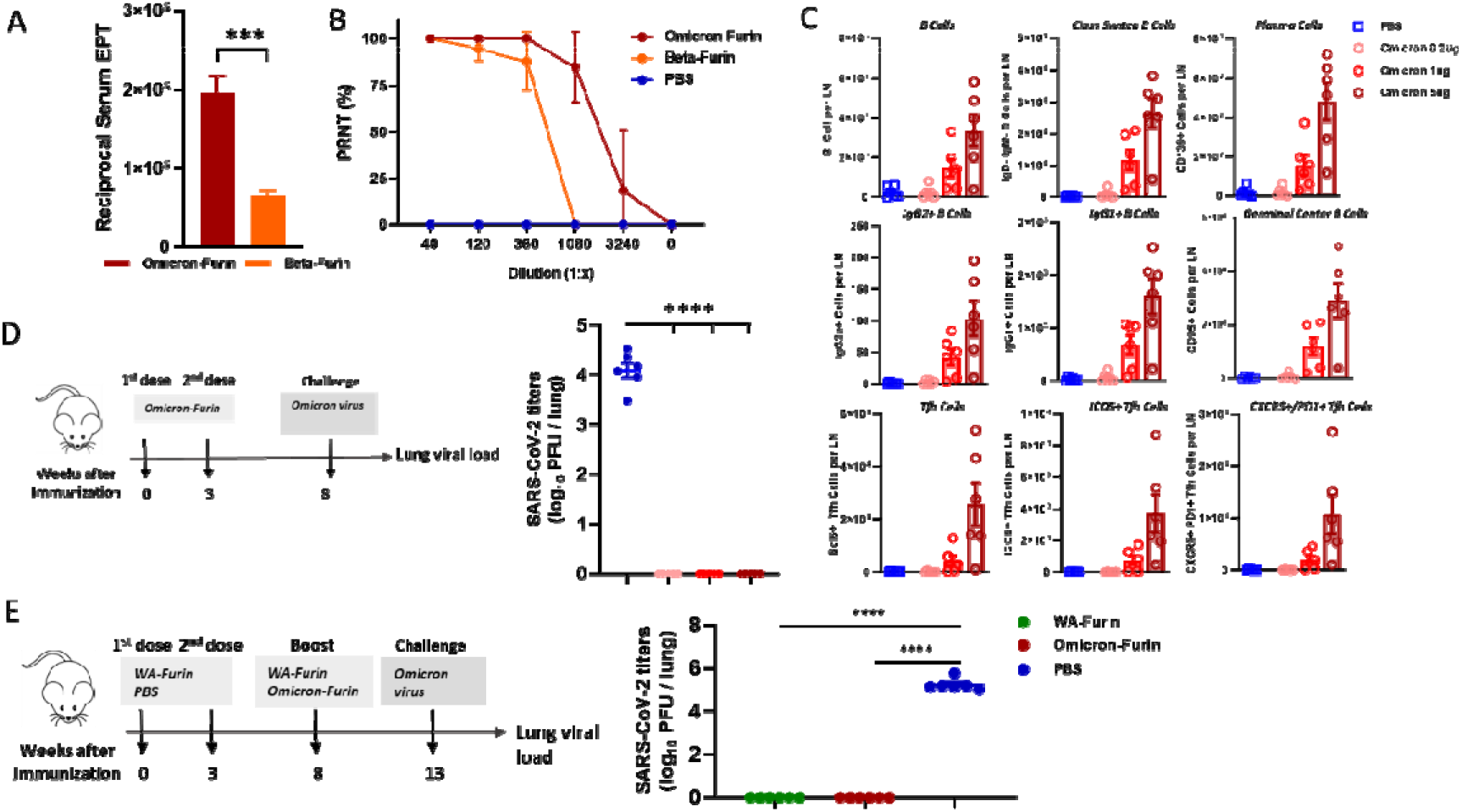
Immunization with Omicron-specific vaccine produces robust protection against Omicron challenge. BALB/c mice were intramuscularly injected with 5 μg Omicron-Furin or Beta-Furin mRNA twice. The Day 14 sera post booster shot were evaluated for total antibodies binding to Omicron S protein by ELISA (**A**) and for nAb responses to live Omicron virus by PRNT (**B**). Plots: mean±SEM; n=6; ***p<0.001 by Student’s t-test. (**C**) BALB/c mice were injected with different doses of Omicron-furin mRNA. Twelve days post injection, the draining lymph nodes (LN) were collected and analyzed by flow cytometry for the abundance of indicated cell types. (**D**) K18 hACE2 mice were immunized with Omicron-furin mRNA as illustrated (left panel). Five weeks after the booster injection, the animals were challenged with the Omicron live virus and the lung virus titer was quantified after four days. (**E**) K18 hACE2 mice were immunized and boosted as illustrated (left panel). Five weeks after booster, the animals were challenged with the Omicron live virus and the lung virus titer was quantified after four days. Plots: mean±SEM; n=6. ****p<0.0001 by one-way ANOVA. EPT: endpoint titer.

To characterize the effect of vaccination on lymphocyte activation and maturation, BALB/c mice were sacrificed twelve days after a single dose of Omicron mRNA at 0.2 ug, 1.0 ug or 5.0 ug, and the draining lymph node (LN) at the injection side was analyzed by flow cytometry. We noticed a dose-dependent increase in total B cells, class-switching B cells, germinal center B cells and plasma cells. Interestingly, both IgG2+ and IgG1+ B cells were increased at a comparable rate, suggesting a balanced T_H_1-T_H_2 response (**Figure 5C**). Furthermore, the vaccination with Omicron mRNA greatly increased the number of total T cells, CD4+ and CD8+ T cells (**Fig. S5**). Remarkably, T follicular helper cells (Tfh), including the ICOS+ and CXCR5/PD1+ subsets, were also dramatically increased by the vaccination, especially at the highest dose, 5 ug (**Figure 5C**), which could contribute to the B cell proliferation and class switching in the germinal center.

To investigate whether Omicron mRNA can indeed elicit strong variant-specific immunity, K18 hACE2 mice were immunized with the mRNA at 0.2 ug, 1.0 ug and 5.0 ug twice at a three-week interval. Five weeks after the booster injection, the animals were challenged with the Omicron strain virus and the virus titer in lungs was quantified four days later. As compared to control animals, no sign of viral replication was detected in Omicron mRNA-vaccinated mice, even at the lowest dose (**Figure 5D**), indicating strong immunity again this specific variant.

As the Omicron strain led to increasing number of breakthrough cases even in people already receiving three-dose vaccination, we wanted to investigate whether using Omicron-Furin mRNA solely as a boosting immunogen could provide enhanced protection. Hence, we set up an *in vivo* challenge study to simulate the real-world scenario (**Figure 5E**). Two doses of WA-Furin mRNA were administrated into K18-ACE2 transgenic mice as described previously. Five weeks after the second dose, the animals received the booster shot of either 5μg WA-Furin mRNA or Omicron-Furin mRNA. After another five weeks, the animals were challenged with live Omicron strain before lung virus loads were quantified. Surprisingly, while the control group displayed high viral titer, up to 5X 10^5 PFU per lung, vaccinated mice showed no detectable viral replication in the lung, suggesting that both WA-1 and Omicron-based booster mRNAs provided substantial and similar protection against Omicron.

### Chimeric RBD-based mRNA vaccine elicits broad protection against Omicron and Delta

With the new omicron mRNA vaccine, we sought to determine whether it could provide sufficient cross-reactive immunity against other VOCs. The ELISA and pseudovirus assays indicated that the Omicron-Furin mRNA vaccine elicited only limited immunity against VOCs, especially Delta (data not shown). Hence, in an effort to generate a vaccine with broad cross-reactivity against other VOCs, we constructed a chimeric VOC immunogen by inserting the RBD domain of the Delta variant directly upstream of the Omicron RBD within the Omicron spike backbone (**Figure 6A**). We first confirmed that the translation product of the chimeric mRNA could still bind to its natural receptor, ACE2 by flow cytometry (**Figure S4**). To evaluate the immunogenicity and efficacy of this chimeric design, mice were immunized twice with LNP-formulated mRNA as described above. The sera were collected two weeks following the second dose and then analyzed for the titers of binding antibodies and nAbs against various VOCs. The ELISA results showed that the chimeric Delta RBD-Omicron mRNA outperformed the original Omicron mRNA in the generation of binding antibodies against WA, Beta and Gamma variants (**Figure 6B**). Surprisingly, although a moderately higher titer against Delta was seen with the chimeric mRNA vaccination, it did not reach statistical significance. As the readout of ELISA assay is not a direct indicator of neutralizing capability, we used the same serum panel to further quantify the nAb titer against Delta and two Omicron pseudoviruses (**Figure 6C**). The IC50 of neutralization showed that compared to Omicron-Furin mRNA, the Delta RBD-Omicron immunization induced similar neutralization activity towards not only the original Omicron (BA.1), but also sublineages including, R346K and the more transmissible BA.2. The absolute IC50 was 5595 with Omicron-Furin and 4945 with Delta RBD-Omicron against the ancestral omicron strain, while the corresponding readout jumped to 8485 and 7589 against Omicron R346K, respectively. Notably, there was a significant increase in the nAb titer against the Delta variant when mice were immunized with the chimeric Delta RBD-Omicron mRNA as compared to mice immunized with the Omicron-Furin mRNA. Taken together, this chimeric design offers a powerful strategy to develop mRNA vaccines with broad protection capacity against COVID-19 and other infectious diseases.

**Figure 6.**
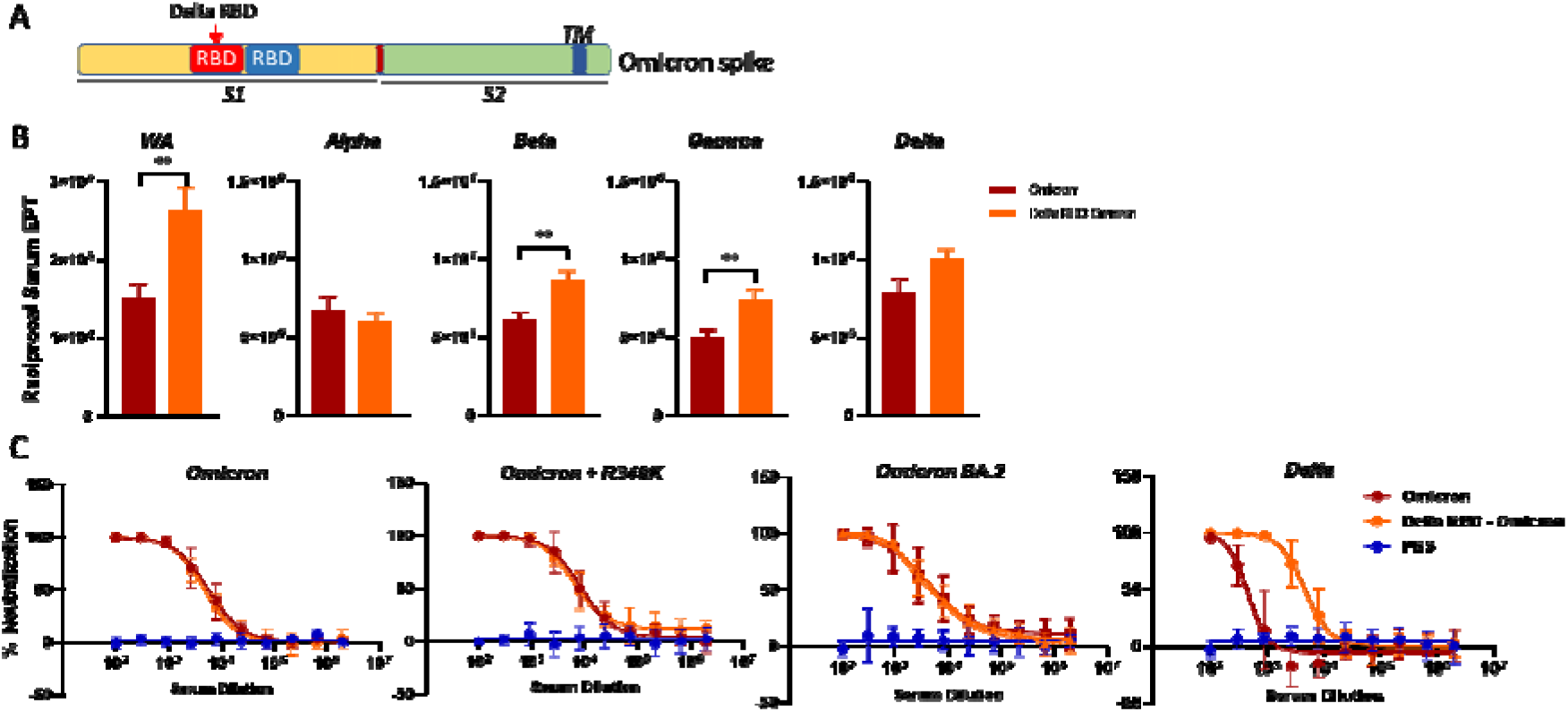
Delta RBD-Omicron immunization induces potent and broad neutralization activity against SARS-COV-2 variants. (**A**) The schematic design of chimeric Delta RBD-Omicron mRNA. BALB/c mice were immunized with the indicated mRNA twice with a 3-week interval. The Day 14 sera post booster were evaluated for total binding antibodies specific to the S proteins of designated VOCs by ELISA (**B**) and for nAb responses to the indicated SARS-CoV-2 variants by pseudovirus assay (**C**). Plots: mean±SEM; n=6. **p<0.01 by Student’s t-test. EPT: endpoint titer.

## Discussion

Prophylactic nucleic acid vaccines can deliver the nucleotide sequence that codes for virus-derived but nonpathogenic proteins into host cells, thus mimicking a native infection to elicit an immune response. Unlike DNA, mRNA vaccines eradicate the need for nucleic acid to enter the nucleus to achieve expression, and they are less likely to be integrated into the host genome. There are currently two mRNA-based SARS-CoV-2 vaccines authorized by FDA and widely disseminated. These vaccines encode for the S protein, the major surface protein on the coronavirus virion responsible for anchoring onto target cells, and thus the predominant virus-encoded target for nAb elicited by natural infection. Although clinical trials and real-world data have affirmed the safety and effectiveness of these FDA-authorized COVID-19 vaccines, more and more breakthrough infections have been reported for predominant VOCs. For example, the effectiveness of BNT162b2 against Delta-caused infection plummeted to 42%, as compared to 95% against the ancestral WA strain^9^, highlighting the need of developing vaccines that can offer wide protection against persistently emerging VOCs.

As each VOC possesses its unique set of mutations in the S protein, this will invariably result in distinct pools of epitopes being presented to lymphocytes by antigen presenting cells (APCs). Hence, we first evaluated the protection conferred following vaccination with mRNA vaccines encoding the S proteins of VOCs that emerged prior to Omicron. Our *in vitro* and *in vivo* data clearly demonstrated the premise that the strongest immunity against individual VOCs can be achieved by vaccination with variant-specific mRNA. For example, immunization with Delta S mRNA provided the best protection against Delta, but not to other VOCs (**Figure 2**). We also noticed that WA and Beta mRNA, especially the latter, provided the widest breadth of coverage for other VOCs. This could be explained by some mutations within Beta S protein, especially in the N-terminal and RBD. These mutations are also present in other VOCs, for example, L18, K417, E484, N501, D614, some of which are known to cause great immune escape ^27^. Interestingly, Beta S mRNA has also been selected by Moderna to test in phase II either as a monovalent antigen or by mixing with mRNA-1273 to tackle the emerging VOCs^28,29^.

With the emergence and rampage of Omicron during the course of our candidate vaccine development, we further tested the effectiveness of non-Omicron VOC mRNA vaccines against this new variant. Not surprisingly, none of these candidates induced strong immunity when challenged with Omicron (**Figure S3**). This could be explained by the more than 30 mutations in the S protein of Omicron, a mutational burden which far exceeded that of preceding VOCs. This observation is consistent with the complete escape of Omicron from most of EUA-approved neutralizing antibody and convalescent sera therapies^30^. To recapitulate our previous observation that VOC-matched vaccine induces the strongest nAb response in a VOC-dependent manner, we constructed a new mRNA vaccine encoding the S protein of Omicron and it indeed provided superior protection against this new VOC (**Figure 5A, B**). The notion was reciprocally confirmed by the observation that Omicron mRNA did not induce strong immunity against other VOCs (**Figure 6**). Recently, a new subvariant of Omicron, BA.2, has quickly spread across the globe and dominated the new cases in many regions, such as Europe, Australia, and China. It is exciting to note that our Omicron mRNA vaccine can elicit strong neutralizing titers against BA.2 (Figure 6C).

As the world is witnessing an astounding number of Omicron-related breakthrough cases even in people have received a third dose of currently available mRNA vaccines, a serious concern has been raised of whether a booster is adequate to stop its spread. Hence, we compared Omicron-specific booster to WA-specific booster to explore any additive protection. However, our data argued that the heterologous booster scheme is not superior to boosting with the ancestral WA mRNA in terms of establishing immunity against Omicron (**Figure 5E**). Similar conclusion was drawn by a research group from NIH, when they challenged mRNA-1273-vaccinated macaques with live omicron virus^31^. A plausible explanation is although Omicron S protein has ~35 mutations, it still exhibits 97% similarity to the ancestral WA strain in the amino acid sequence. As a result, most of the epitopes presented to lymphocytes could remain the same, and the consequence of antigenic drift contributed by those ~35 mutations could be masked by the surge of nAbs generated after the third dose. Hence, individuals with prior immunity from vaccination may not necessarily benefit from a change in vaccinating antigens. Moreover, considering the cost and time needed to put variant specific mRNA vaccine to practical use, the homologous boost scheme remains a scientifically proven and economically feasible option at hand in the fight against COVID-19.

Nonetheless, the concept of universal vaccine is still very appealing for at least two reasons. One is the virus could keep accumulating more mutations to eventually nullify the effectiveness of existing mRNA vaccines. The other is the idea can be applied to establish immunity against viruses causing different diseases. For example, dimeric RBDs have been ligated in tandem to target MERS, SARS and COVID-19^32^. In comparison, our chimeric design included the incorporation of the RBD of Delta variant in the Omicron spike mRNA which offers a larger pool of epitopes. Remarkably, we found that the resultant mRNA restored the strong protection against Delta infection while retaining the effective immunity against Omicron (**Figure 6**). Similar strategies have also been explored to target the *Sarbecovirus* subgenus with a single-molecule antigen^33^, suggesting broad applications of the chimeric vaccine design to deliver effective protection against a wide panel of diseases.

Another unique feature of our mRNA vaccine is the mutation of furin cleavage site between the S1 and S2 domains of S. This cleavage is believed to have emerged during viral transmission from its zoonotic host to humans and is one of the key attributes to explain the high transmissibility of SARS-CoV-2 in humans^34^. The mutation is mainly to address the concern that circulating S1 was detected in the plasma of vaccinated subjects^35^. Although the clinical consequence of the free S1 moiety has not been full established, studies have suggested that S1 can be taken up by many critical organs, such as liver, kidney, spleen, and even cross the blood-brain barrier to gain access to the brain. As S1 contains the intact RBD, it could still bind to ACE2 and trigger the downstream signaling events that may lead to inflammation and lung damage^36,37^. Hence, we mutated the furin cleavage site and confirmed the abrogation of S1 liberation in both the cell lysates and conditioned medium. Although a similar design was adopted in the recombinant protein-based NVX-CoV2373 by Novovax, it is not present in the two mRNA-based Covid-19 vaccines authorized by FDA. Another potential benefit of furin cleavage mutation is by retaining the full-length S protein within the cell and on the cell surface (**Figure 1B-D**), a larger pool of antigens could become available for presentation to induce the adaptive immunity. Indeed, S protein with furin cleavage mutation even binds with higher affinity to ACE2^38^.

Taken together, our in-house designed mRNA vaccine represents a potentially safer alternative to existing products on the market and can induce stronger immunity against prevalent VOCs, including Omicron and Delta. Our chimeric design will also facilitate the development of next-generation vaccines that achieve the balance between effectiveness and coverage, not only for the variants of SARS-CoV-2 but also for other viruses.

## Materials and Methods

### In vitro transcription and purification of RNA

To generate the template for RNA synthesis, the sequences of the SARS-Cov-2 Spike protein of VOC were codon optimized and cloned into pVAX1-based backbone which features 5⍰-UTR, 3⍰-UTR and Poly-A tail. To increase the protein stability, 2P mutations at positions 986-987 were introduced. The plasmid DNA was produced in bacteria, purified and linearized by a single-site restriction enzyme digestion. The template DNA was purified, spectrophotometrically quantified, and in vitro transcribed by T7 RNA polymerase (Cat: M0251, NEB) in the presence of a trinucleotide cap1 analogue, m7(3OMeG)(5⍰)ppp(5⍰)(2OMeA)pG (Cat: N-7113, TriLink), and of N1-methylpseudouridine-5’-triphosphate (Cat: N-1081, TriLink) in place of uridine-5’-triphosphate (UTP). After the reaction, DNase I (Cat: M0303, NEB) was added to remove the template DNA and the mRNA was purified by LiCl precipitation (Cat: AM9480, ThermoFisher).

### mRNA formulation

LNPs were prepared by microfluidic mixing a buffered solution of mRNA with an ethanol solution of lipids [distearoylphosphatidylcholine (DSPC), cholesterol, 1,2-Dimyristoyl-rac-glycero-3-methoxypolyethylene glycol-2000 (DMG-PEG2000), and ionizable lipid. The LNPs were concentrated by dialysis against an aqueous buffer system, following a 0.2 μm sterile filtration. The LNPs were tested for mRNA concentration, encapsulation efficiency, particle size, pH, and osmolality.

### In vitro mRNA expression

Monocytes were isolated from PBMCs and differentiated into DCs in presence of GM-CSF (Cat: 300-03, Peprotech) and IL-4 (Cat: 200-04, Peprotech). Between day 6-day 8, cells were transfected with mRNA by the NeonTM electroporation transfection system (Cat: MPK5000, ThermoFisher). 24 hours post-transfection, the cells were collected and subjected to flow cytometry as described below to check the expression of spike. With 293T adherent cells, mRNA (2.5ug) of WT vs Mutant from five variants (Washington, Alpha, Beta, Gamma and Delta) were transfected with Lipofectamine™ MessengerMAX™ Transfection Reagent (2ul) and cultured for 72 hours at 37oC using 293T adherent cells with 0.5ml of DMEM media with 10% FBS in each well of a 24-well cell culture-treated plate. Transfected cells from each well were dislodged with 400ul of TrypLE at 72 hours and neutralized with its own media. Cell pellets were collected after spinning down at 550g for 2 minutes by removing supernatant for each well.

### Flow cytometry

The collected cell pellets were washed with 250ul of FACS buffer (DPBS + 0.5% BSA) in 96 wells treated plate followed by 30-minute incubation with the in-house STI-2020 (for DCs) or 10A3 (for 293T cells) primary antibody (1:1000) to detect SARS Cov-2 spike. 200ul FACS buffer was used to wash the cells twice with the same speed and time after 30-minute incubation, followed by rat anti-human Fc antibody conjugated to APC (Cat: 410712, BioLegend; 1:100 dilution) for 15 min on ice in the dark for secondary detection. The cells were spun down and the pellets were washed twice with the same speed and time of centrifugation using 200ul FACS buffer and resuspended in 200ul FACS buffer. The fluorescent intensity of positive cells within the gated population was detected by the Attune NxT Flow Cytometer (ThermoFisher) using 100ul of acquisition volume setting.

### SARS-CoV-2 Virus

SARS-COV-2 viruses were obtained from BEI resources (Washington strain NR-52281; Alpha variant NR-54000; Beta Variant NR-54009; Gamma variant NR-54982; Delta variant NR-55611 or NR-55672; Lambda variant NR-55654 and Omicron NR-56461). VeroE6 monolayers were infected at an MOI of 0.01 in 5ml virus infection media (DMEM + 2% FCS + 1X Pen/Strep). Tissue culture flasks were incubated at 36°C and slowly shaken every 15 minutes for a 90-minute period. Cell growth media (35mL) was added to each flask and infected cultures were incubated at 36°C/5% CO2 for 48 hours. Media was then harvested and clarified to remove large cellular debris by room temperature centrifugation at 3000 rpm.

### Animals and in vivo studies

7-week-old BALB/cJ female mice were purchased from the Jackson Laboratory. All protocols were approved by the Institutional Animal Care and Use Committee (IACUC). mRNA formulations were diluted in 50 uL of 1X PBS, and mice were inoculated IM into the same hind leg for both prime and boost. There was 3 weeks interval between prime and boost. Two weeks after boost, mice blood was collected from retro-orbital for ELISA and pseudovirus neutralization assay.

### ELISA

Ni-NTA HisSorb plates (Qiagen) were coated with 50ng/well of S1 proteins (all from Sino Biological, Cat: 40591-V08H, 40589-V08B6, 40589-V08B7, 40589-V08B8, 40589-V08B16) in 1X PBS at 4°C overnight. To block the plate, Blocker Casein (Cat: 37528 Thermo) was used for 1 hour at room temperature (RT). After standard washes and blocks, plates were incubated with serial dilutions of sera for 1 hour at RT. Following washes, goat anti-mouse IgG (H+L)-HRP conjugate (Cat: #1721011, Bio-Rad) were used as secondary Abs, and Pierce TMB substrate kit (Cat: 34021, Thermo Fisher) was used as the substrate. The absorbance was measured at 450 nanometers using a BioTek Cytaktion 5 plate reader. Endpoint tiers were calculated as the dilution that emitted an optical density exceeding 4X background (secondary Ab alone).

### T follicular helper (Tfh) and B cell phenotype detection in lymph node by flow cytometer

For mouse T cell and B cell analysis in lymphoid tissues, lymph node cells were stained for viability and extracellular antigens with directly labelled antibodies, CD95 (Biolegend 152606), IgD (Biolegend 405710), GL7 (Biolegend 144617), Live/Dead Fixable Yellow Dead Cell Staining (Thermo Scientific L34968), CD19 (Biolegend 115555), Biotinylated SARS-CoV-2-S1 Protein (Biolegend 793804) Anti-Biotin (409003), CD3 (Biolegend 100220), CD3 (Biolegend 100204), CD4 (Biolegend 100540), ICOS (117420), CD8 (Biolegend 557654), CD45 (Biolegend 103134), CXCR5 (Biolegend 145504), PD-1 (Biolegend 135231), CD38 (102712), IgG1 (Biolegend 406620), IgG2a (Biolegend 407114). Cells were washed, fixed and permeabilized by using Ebioscience Foxp3/Transcription Factor Staining Buffer Set (Thermo Scientific 00-5523-00) kit according to manufacture instructions. Permeabilized cells were intracellularly stained with Bcl-6 (Biolegend 358512). Cells were acquired on Attune NxT Flow Cytometer and analyzed with Attune NxT software v 4.2.0.

### Quantify SARS-CoV-2 S-specific T cells in mice

Five weeks after booster, mouse splenocyte were isolated and incubated for 6 hours at 37°C with BD GolgiPlug (BD Bioscience 555028) and with or without spike peptide pools (JPT Peptide Technologies PM-WCPV-S-1). Cells were washed, stained, and analyzed as described in the section above “T follicular helper (Tfh) and B cell phenotype detection in lymph node by flow cytometer”. Antibodies for extracellular antigens are CD3 (BioLegend 100220), CD4 (BioLegend 100540), CD8 (BD Bioscience 557654), CD44 (BioLegend 103040), anti-I-A/I-E (BioLegend 107608), and Live/Dead Fixable Yellow Dead Cell Staining (Thermo Scientific L34968), and for intracellular antigens are TNF-α (BioLegend 506304) and IFN-γ (BioLegend 505810).

### Pseudovirus neutralization assay

SARS-CoV-2 Spike pseudotyped ΔG-VSV-luciferase was generated by nucleofection of BHK cells (maintained in DMEM/F12 with 10%FBS and 5%TPB) with Spike-expressing plasmid followed by transduction with G-pseudotyped ΔG-luciferase (G*ΔG-luciferase) rVSV (Kerafast) 18-24 hours later. The supernatant containing pseudovirus was collected following 24 hours and stored at −80°C. Pseudovirus was normalized for luciferase expression using G*ΔG-luciferase VSV of known titer as the standard. For neutralization testing, HEK-Blue 293 hACE2-TMPRSS2 cells (Invivogen; maintained in DMEM with 10% FBS) were plated to white-walled 96-well plates at 40,000 cells/well and incubated at 37°C/5% CO2. The next day, SARS-CoV-2 Spike pseudotyped ΔG-VSV-luciferase was incubated with a dilution series of mouse serum (dilutions as indicated) and anti-VSV-G (Kerafast; 1 μg/mL) antibody for 30 minutes at room temperature and added to the HEK-Blue 293 hACE2-TMPRSS2 cells. Transduced cells were incubated for 24 hours at 37°C/5% CO2 and luminescence measured by addition of 40μl of ONE-Glo reagent (Promega) with detection using a Tecan Spark plate reader. The percent inhibition was calculated using the formula 1-([luminescence of serum treated sample]/[average luminescence of untreated samples] × 100. The average of quadruplicate samples was included in the analyses.

### Plaque Reduction Neutralization Test (PRNT)

Simian VeroE6 cells were plated at 18×103 cells/well in a flat bottom 96-well plate in a volume of 200 μl/well. After 24 hours, a serial dilution of seropositive blood serum is prepared in a 100 μl/well at twice the final concentration desired and live virus was added at 1,000 PFU/100μl of SARS-CoV-2 and subsequently incubated for 1 hour at 37°C in a total volume of 200 μl/well. Cell culture media was removed from cells and sera/virus premix was added to VeroE6 cells at 100 μl/well and incubated for 1 hour at 37°C. After incubation, 100 μl of “overlay” (1:1 of 2% methylcellulose (Sigma) and culture media) is added to each well and incubation commenced for 3 days at 37°C. Plaque staining using Crystal Violet (Sigma) was performed upon 30 min of fixing the cells with 4% paraformaldehyde (Sigma) diluted in PBS. Plaques were assessed using a light microscope (Keyence).

### Challenge study

K18-hACE2 transgenic mice were purchased from Jackson laboratory and maintained in pathogen-free conditions and handling conforms to the requirements of the National Institutes of Health and the Scripps Research Institute Animal Research Committee. 8-12 weeks old mice were injected with the indicated administration technique under isoflurane anesthesia in the right hind flank area for IM injections. Mice were infected intranasally with 10000 PFU of SARS-CoV-2 in total volume 50 μL.

### Plaque assay

VeroE6 cells were plated at 3×10e5 cells/well in 24 well plates in volume 400 μl/well. After 24 h. medium is removed, and serial dilution of homogenized lungs were added to Vero cells and subsequently incubated for 1 h at 37°C. After incubation, an overlay (1:1 of 2% methylcellulose (Sigma) and culture media) is added to each well and incubation commenced for 3 d at 37°C. Plaque staining was performed using Crystal Violet as mentioned above.

### Statistics

Statistical significance of differences between experimental groups was determined with Prism software (Graphpad). All data are expressed as standard error mean (SEM). ****P < 0.0001, ***P < 0.001, **P < 0.01, and *P < 0.05 by unpaired two-tailed t tests or one- or two-way analysis of variance (ANOVA).

## Author Contributions

R.C., H.J., Q.H., Y.Z., N.S., X.L. conceptualized and designed experiments. Q.H., Y.Z., N.S., X.L., H.S., V.C., Y.K., J.X., A.F., K.S., Z.W., Z.W., Y.H., Y.Z., L.K., C.Z., H.W. performed experiments. Q.H., Y.Z., N.S., X.L., H.S., H.P., C.P., R.A., H.X., R.C. analyzed and interpreted the data. Q.H., Y.Z., N.S., X.L. and R.C. wrote the paper.

## Competing interests

Sorrento authors own options and/or stock of the company. This work has been described in one or more provisional patent applications. H.J. is an officer at Sorrento Therapeutics, Inc.

## Acknowledgments

We thank Lisa Kerwin, Nancy Du and Yanliang Zhang for providing reagents; Avery Coyle and Charlotte Sadaka for helpful discussions; Brian Sun, Mike Ruse, Jr., Elizabeth Orr, Gali Steinberg-Tatman and Maggy Smith for patent applications.

---

**Figure S1.**
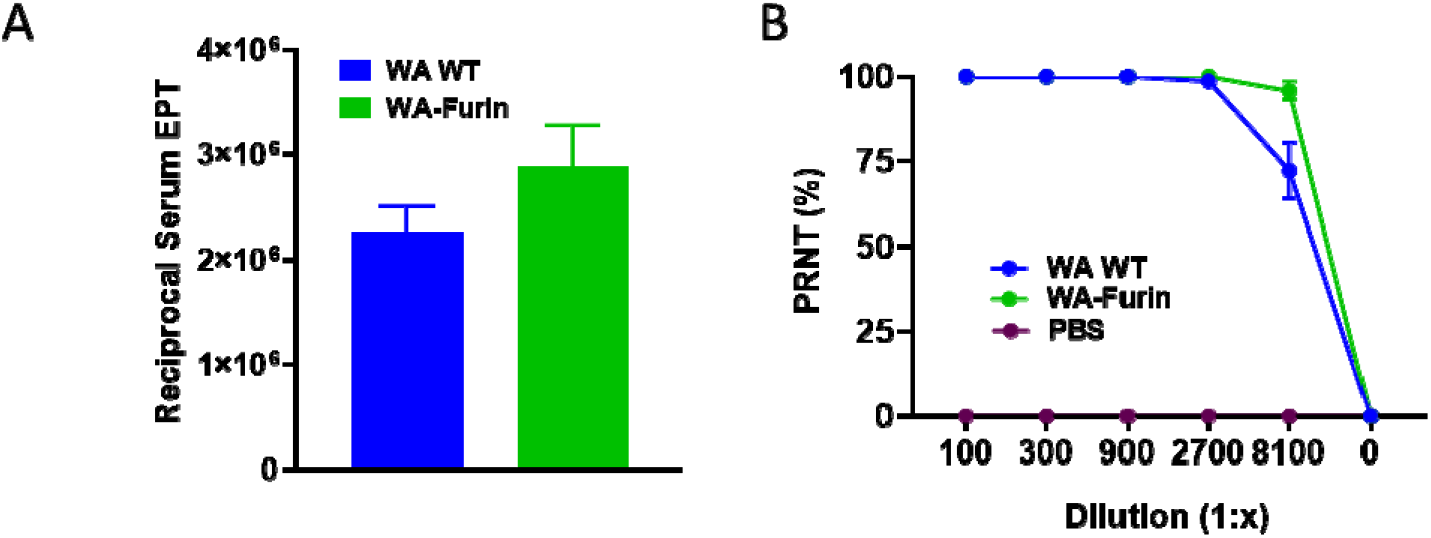
Evaluation of wild-type and Furin cleavage mutant mRNA *in vivo*. BALB/c mice were immunized twice with either wild-type or furin cleavage mutant mRNA encoding the S protein of the ancestral WA strain. The day 14 sera after booster were analyzed for the titers of total binding antibodies by ELISA (**A**) and for those of neutralizing antibodies by PRNT (**B**). Plots: mean±SEM; n=6.

**Figure S2.**
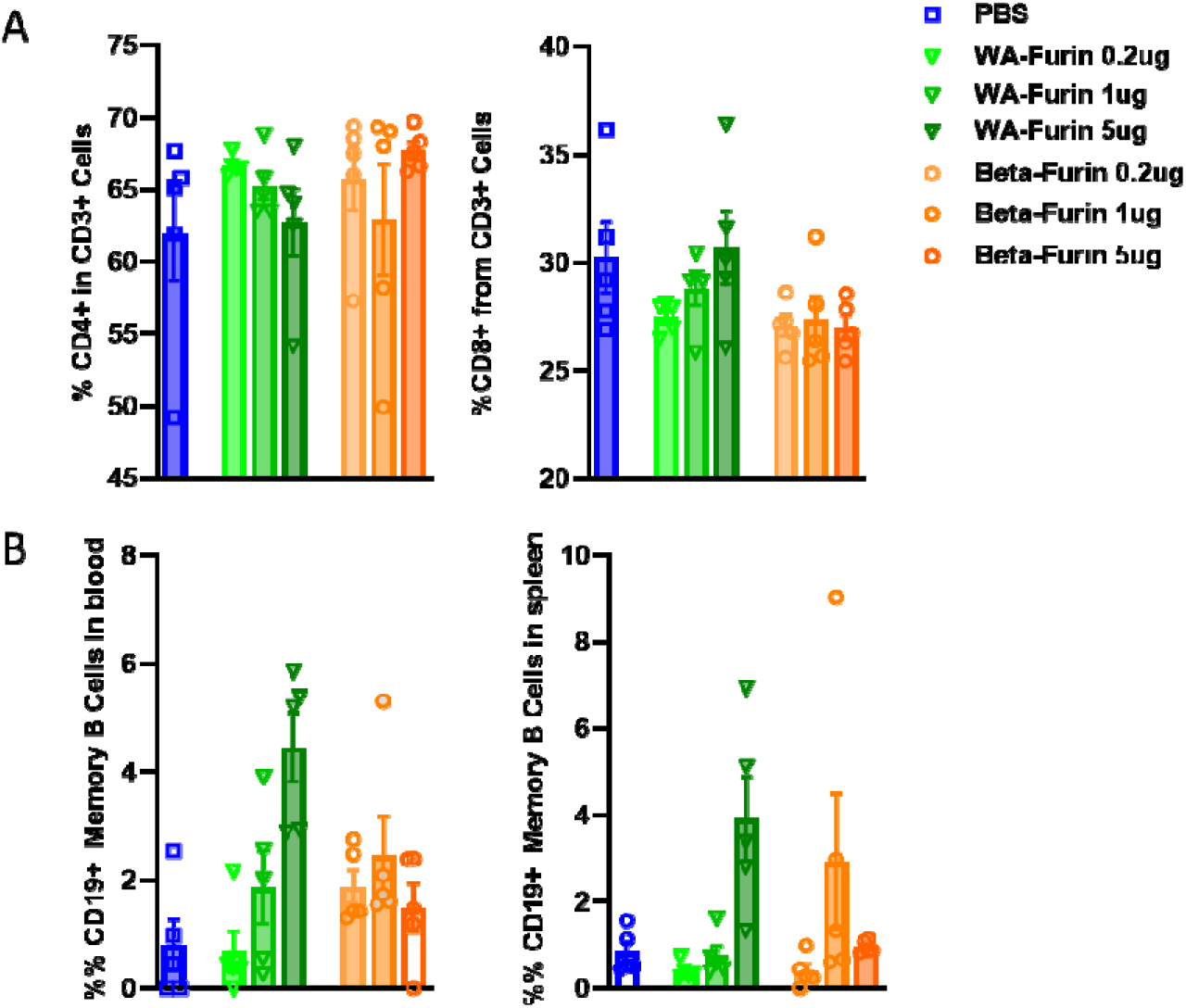
Phenotyping of T cell and B cells from blood and spleen in immunized mice. BALB/c mice were immunized twice with WA-Furin or Beta-Furin mRNA at indicated doses. Seven weeks after booster, CD4+ T cells, CD8+ T cells from spleen (**A**), and memory B cells from spleen and blood (**B**) were quantified by flow cytometry. Bars: mean±SEM; n=5.

**Figure S3.**
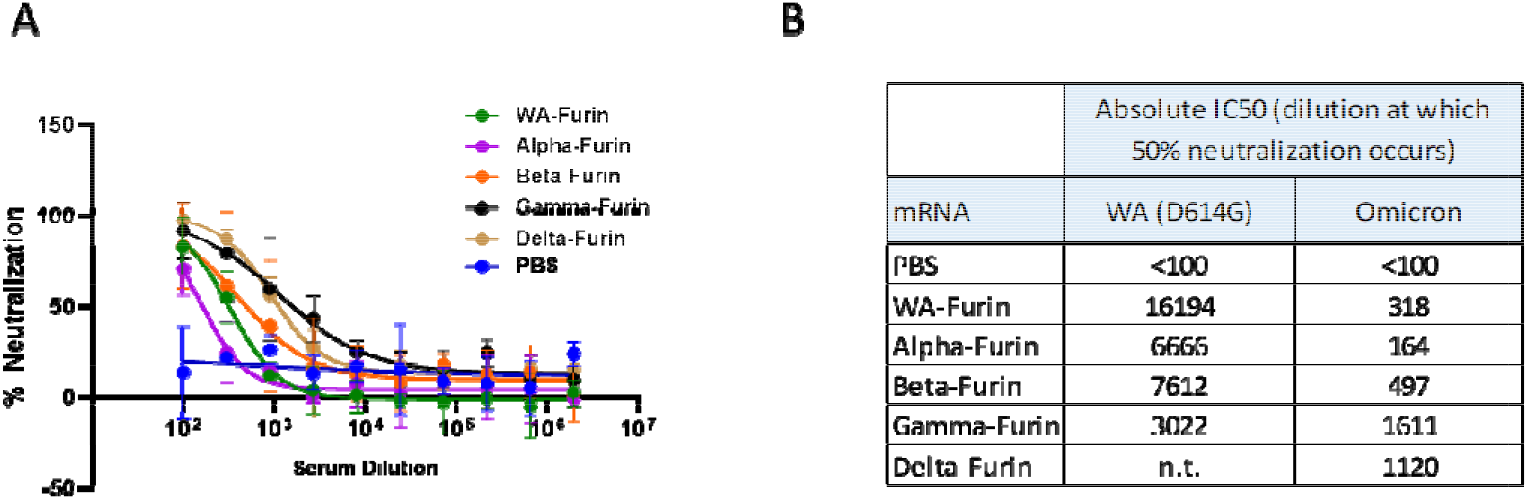
Quantification of neutralizing antibodies against Omicron pseudovirus. BALB/c mice were immunized with furin cleavage mutant mRNAs of indicated VOCs. (**A**) The Day 14 sera after booster shot were evaluated for nAb responses against Omicron by pseudovirus assay. Plots: mean±SEM; n=6. (**B**) Summary of average IC50 dilution of the neutralization assay.

**Figure S4.**
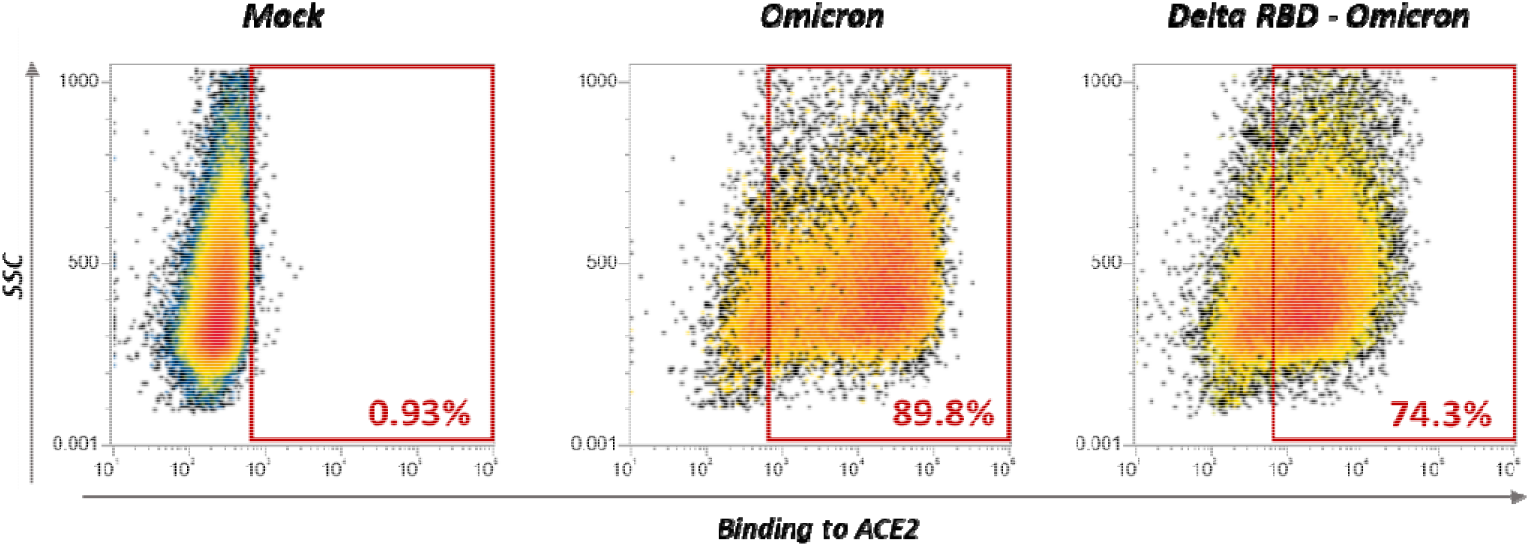
Expression of Omicron and Delta RBD-Omicron mRNA. 293T cells in 24-well plate were transfected with 1ug indicated mRNA, and collected 24 hours post-transfection. The flow cytometry showed the surface expression of spike protein in transfected 293T cells stained with recombinant human ACE2 protein.

**Figure S5.**
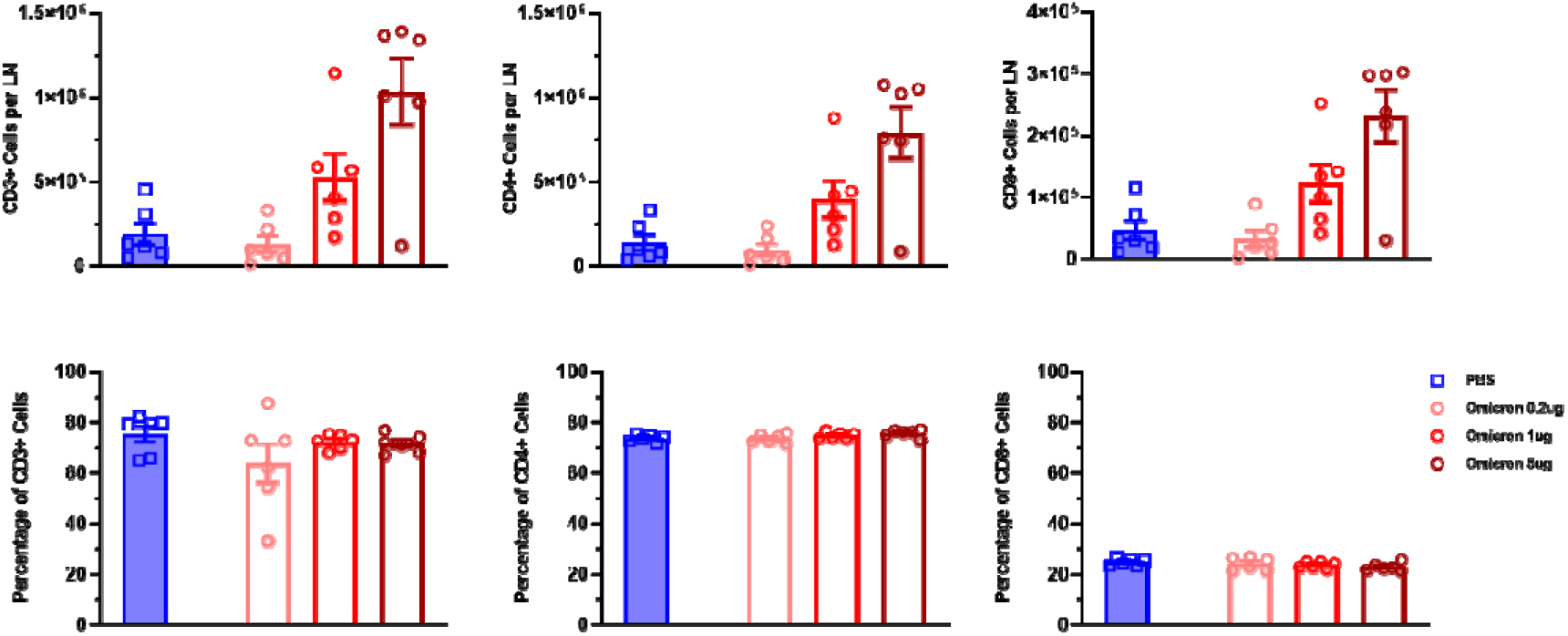
Omicron mRNA vaccination promotes T lymphocyte proliferation in draining LNs. BALB/c mice were injected with Omicron-furin mRNA at indicated doses. Twelve days post vaccination, the draining LNs were collected and analyzed by flow cytometry to determine the abundance of T lymphocytes. LN: lymph node. Bars: mean±SEM; n=6.

## Notes

### Competing Interest Statement

The authors have declared no competing interest.

## Reference

1. Adam, D. The pandemic’s true death toll: millions more than official counts. Nature 601, 312–315 (2022).

2. WHO – COVID19 Vaccine Tracker. https://covid19.trackvaccines.org/agency/who/.

3. Otto, S. P. et al. The origins and potential future of SARS-CoV-2 variants of concern in the evolving COVID-19 pandemic. Curr. Biol. 31, R918–R929 (2021).

4. Liu, Z. et al. Identification of SARS-CoV-2 spike mutations that attenuate monoclonal and serum antibody neutralization. Cell Host Microbe 29, 477–488.e4 (2021).

5. Abu-Raddad, L. J., Chemaitelly, H. & Butt, A. A. Effectiveness of the BNT162b2 Covid-19 Vaccine against the B.1.1.7 and B.1.351 Variants. N. Engl. J. Med. 385, 187–189 (2021).

6. Planas, D. et al. Reduced sensitivity of SARS-CoV-2 variant Delta to antibody neutralization. Nature 596, 276–280 (2021).

7. Mlcochova, P. et al. SARS-CoV-2 B.1.617.2 Delta variant replication and immune evasion. Nature 599, 114–119 (2021).

8. Farinholt, T. et al. Transmission event of SARS-CoV-2 Delta variant reveals multiple vaccine breakthrough infections. medRxiv 2021.06.28.21258780 (2021) doi:10.1101/2021.06.28.21258780.

9. Puranik, A. et al. Comparison of two highly-effective mRNA vaccines for COVID-19 during periods of Alpha and Delta variant prevalence. MedRxiv Prepr. Serv. Health Sci. 2021.08.06.21261707 (2021) doi:10.1101/2021.08.06.21261707.

10. Xu, Z., Liu, K. & Gao, G. F. Omicron variant of SARS-CoV-2 imposes a new challenge for the global public health. Biosaf. Health (2022) doi:10.1016/j.bsheal.2022.01.002.

11. Cao, Y. et al. Omicron escapes the majority of existing SARS-CoV-2 neutralizing antibodies. 2021.12.07.470392 (2021) doi:10.1101/2021.12.07.470392.

12. Lee, I.-J. et al. Omicron-specific mRNA vaccine induced potent neutralizing antibody against Omicron but not other SARS-CoV-2 variants. 2022.01.31.478406 (2022) doi:10.1101/2022.01.31.478406.

13. Zhao, X. et al. Reduced sera neutralization to Omicron SARS-CoV-2 by both inactivated and protein subunit vaccines and the convalescents. 2021.12.16.472391 (2021) doi:10.1101/2021.12.16.472391.

14. Wu, L. et al. SARS-CoV-2 Omicron RBD shows weaker binding affinity than the currently dominant Delta variant to human ACE2. Signal Transduct. Target. Ther. 7, 1–3 (2022).

15. Shang, J. et al. Cell entry mechanisms of SARS-CoV-2. Proc. Natl. Acad. Sci. 117, 11727–11734 (2020).

16. Francis, D. M. et al. Directing an mRNA-LNP vaccine toward lymph nodes improves humoral and cellular immunity against SARS-CoV-2. 2021.08.25.457699 (2021) doi:10.1101/2021.08.25.457699.

17. Peacock, T. P. et al. The furin cleavage site in the SARS-CoV-2 spike protein is required for transmission in ferrets. Nat. Microbiol. 6, 899–909 (2021).

18. Ogata, A. F. et al. Circulating Severe Acute Respiratory Syndrome Coronavirus 2 (SARS-CoV-2) Vaccine Antigen Detected in the Plasma of mRNA-1273 Vaccine Recipients. Clin. Infect. Dis. 74, 715–718 (2022).

19. Nyiro, J. U. et al. Agreement between ELISA and plaque reduction neutralisation assay in Detection of respiratory syncytial virus specific antibodies in a birth Cohort from Kilifi, coastal Kenya. Wellcome Open Res. 4, 33 (2019).

20. Schmidt, N. J., Dennis, J. & Lennette, E. H. Plaque reduction neutralization test for human cytomegalovirus based upon enhanced uptake of neutral red by virus-infected cells. J. Clin. Microbiol. 4, 61–66 (1976).

21. Radvak, P. et al. B.1.1.7 and B.1.351 variants are highly virulent in K18-ACE2 transgenic mice and show different pathogenic patterns from early SARS-CoV-2 strains. 2021.06.05.447221 (2021) doi:10.1101/2021.06.05.447221.

22. Arce, V. M. & Costoya, J. A. SARS-CoV-2 infection in K18-ACE2 transgenic mice replicates human pulmonary disease in COVID-19. Cell. Mol. Immunol. 18, 513–514 (2021).

23. Dong, W. et al. The K18-Human ACE2 Transgenic Mouse Model Recapitulates Non-severe and Severe COVID-19 in Response to an Infectious Dose of the SARS-CoV-2 Virus. J. Virol. (2021) doi:10.1128/JVI.00964-21.

24. Winkler, E. S. et al. SARS-CoV-2 infection of human ACE2-transgenic mice causes severe lung inflammation and impaired function. Nat. Immunol. 21, 1327–1335 (2020).

25. Planas, D. et al. Considerable escape of SARS-CoV-2 Omicron to antibody neutralization. Nature 602, 671–675 (2022).

26. Muik, A. et al. Neutralization of SARS-CoV-2 Omicron by BNT162b2 mRNA vaccine–elicited human sera. Science 375, 678–680 (2022).

27. Winger, A. & Caspari, T. The Spike of Concern—The Novel Variants of SARS-CoV-2. Viruses 13, 1002 (2021).

28. Pajon, R. et al. SARS-CoV-2 Omicron Variant Neutralization after mRNA-1273 Booster Vaccination. N. Engl. J. Med. NEJMc2119912 (2022) doi:10.1056/NEJMc2119912.

29. Waltz, E. COVID vaccine makers brace for a variant worse than Delta. Nature 598, 552–553 (2021).

30. Liu, L. et al. Striking antibody evasion manifested by the Omicron variant of SARS-CoV-2. Nature 602, 676–681 (2022).

31. Gagne, M. et al. mRNA-1273 or mRNA-Omicron boost in vaccinated macaques elicits comparable B cell expansion, neutralizing antibodies and protection against Omicron. 2022.02.03.479037 (2022) doi:10.1101/2022.02.03.479037.

32. Dai, L. et al. A Universal Design of Betacoronavirus Vaccines against COVID-19, MERS, and SARS. Cell 182, 722–733.e11 (2020).

33. Martinez, D. R. et al. Chimeric spike mRNA vaccines protect against Sarbecovirus challenge in mice. Science 373, 991–998 (2021).

34. Whittaker, G. R. SARS-CoV-2 spike and its adaptable furin cleavage site. Lancet Microbe 2, e488–e489 (2021).

35. Ogata, A. F. et al. Circulating SARS-CoV-2 Vaccine Antigen Detected in the Plasma of mRNA-1273 Vaccine Recipients. Clin. Infect. Dis. Off. Publ. Infect. Dis. Soc. Am. ciab465 (2021) doi:10.1093/cid/ciab465.

36. Letarov, A. V., Babenko, V. V. & Kulikov, E. E. Free SARS-CoV-2 Spike Protein S1 Particles May Play a Role in the Pathogenesis of COVID-19 Infection. Biochem. Biokhimiia 86, 257–261 (2021).

37. Suzuki, Y. J. & Gychka, S. G. SARS-CoV-2 Spike Protein Elicits Cell Signaling in Human Host Cells: Implications for Possible Consequences of COVID-19 Vaccines. Vaccines 9, 36 (2021).

38. Laczkó, D. et al. A Single Immunization with Nucleoside-Modified mRNA Vaccines Elicits Strong Cellular and Humoral Immune Responses against SARS-CoV-2 in Mice. Immunity 53, 724–732.e7 (2020).

